# Are Machines-learning Methods More Efficient than Humans in Triaging Literature for Systematic Reviews?

**DOI:** 10.1101/2021.09.30.462652

**Authors:** Seye Abogunrin, Luisa Queiros, Mateusz Bednarski, Marc Sumner, David Baehrens, Andreas Witzmann

**Affiliations:** F. Hoffmann La Roche, Basel, Switzerland; Roche Polska Sp. z o.o., Warsaw, Poland; Averbis GmbH, Freiburg, Germany

**Keywords:** Advanced analytic techniques, Deep learning, Natural language processing, Support vector machine

## Abstract

Systematic literature reviews provide rigorous assessments of clinical, cost-effectiveness, and humanistic data. Accordingly, there is a growing trend worldwide among healthcare agencies and decision-makers to require them in order to make informed decisions. Because these reviews are labor-intensive and time consuming, we applied advanced analytic methods (AAM) to determine if machine learning methods could classify abstracts as well as humans. Literature searches were run for metastatic non-small cell lung cancer treatments (mNSCLC) and metastatic castration-resistant prostate cancer (mCRPC). Records were reviewed by humans and two AAMs. AAM-1 involved a pre-trained data-mining model specialized in biomedical literature, and AAM-2 was based on support vector machine algorithms. The AAMs assigned an accept/reject status, with reasons for exclusion. Automatic results were compared to those of humans. For mNSCLC, 5820 records were processed by humans and 440 (8%) records were accepted and the remaining items rejected. AAM-1 correctly accepted 6% of records and correctly excluded 79%. AAM-2 correctly accepted 6% of records and correctly excluded 82%. The review was completed by AAM-1 or AAM-2 in 52 hours, compared to 196 hours for humans. Work saved was estimated to be 76% and 79% by AAM-1 and AAM-2, respectively. For mCRPC, 2434 records were processed by humans and 26% of these were accepted and 74% rejected. AAM-1 correctly accepted 23% of records and rejected 62%. AAM-2 correctly accepted 20% of records and rejected 66%. The review was completed by AAM-1, AAM-2, and humans in 25, 25 and 85 hours, respectively. Work saved was estimated to be 61% and 68% by AAM-1 and AAM-2, respectively. AAMs can markedly reduce the time required for searching and triaging records during a systematic review. Methods similar to AAMs should be assessed in future research for how consistent their performances are in SLRs of economic, epidemiological and humanistic evidence.

## Introduction

A systematic literature review (SLR) is a specialized type of literature review that is designed to address a specific research question using rigorous, reproducible, and transparent methods. An SLR may focus on various topics, such as clinical trials for a given drug and/or indication, economic evaluation of healthcare technologies, epidemiological information, or humanistic data. Because SLRs are rigorous, there is a growing trend worldwide among health technology agencies (HTAs) and other healthcare decision-makers to require SLRs in order to make informed decisions [1, 2]. In addition, the importance of SLRs in supporting modern evidence-based medicine is shown by the 27-fold increase in the annual rate of published SLRs to 28,959 [3]. However, an SLR is highly time and labor-intensive. An analysis of 195 SLRs in the PROSPERO registry found that it took an average of 67.3 (±31.0) weeks to complete a review through publication [4]. Currently, there are more than 32 million citations cataloged in PubMed, and the list grows by more than 3000 articles daily; thus, generating an SLR requires searching through substantially large numbers of references [5].

Because SLRs are generally meant to contain the most recently available studies, they need to be updated periodically. Moreover, for both novel and updates to SLRs, the task of identifying, examining the text, and triaging the results into accepted and rejected articles for inclusion in the reviews is particularly time-consuming [6, 7]. Automating these tasks could speed up the process as well as free up investigators’ time to tend to other matters. This is why several articles have been investigating how automation can be implemented in the SLR process [8, 9]. A potential solution is to use artificial intelligence (AI), including natural language processing (NLP), in which computer science, and linguistics are used to teach machines to understand human languages and to learn to classify or categorize text [10]. With this method, algorithms can identify and extract unstructured language elements and render them into a form that a machine can understand, following which the text elements can be used for categorization [7].

One method explored is deep learning, a subset of AI [11–13]. It is an advanced analytics technique that teaches a machine to perform a task through learning by example. However, deep learning requires considerable computing power, and it is not always practical to develop a deep learning model from scratch. A common method to deal with this issue is transfer learning. Instead of training a deep model from blank, an already trained model in a similar domain is used to initialize the model. It allows the target model to be trained faster and achieve better results. In terms of NLP, the base model is trained through the use of a very public corpus, for example, all Wikipedia articles, using an axillary task. This allows for grabbing general grammar and language structures, but not texts specific to a specialized field. In this way, a model learns what a noun, adjective, verb is, and so on. This step is called pre-training.

Once a large pre-trained model is established and becomes available for other to use, it can be transferred onto a more specialized model by combining a selected pre-trained model with a classifier (e.g. feed-forward neural network), trained on a smaller dataset, resulting in substantial time savings for the more specialized model. This step is called fine-tuning. During this phase there is no need to use such a big amount of data and computing power - it can be fine-tuned using only target task data, resulting in a classification model ready to be used.

For example, BERT and other Transformer encoder architectures have been very successful on a variety of tasks in natural language processing. They compute vector-space representations of natural language that are suitable for use in deep learning models. The BERT family of models uses the Transformer encoder architecture to process each token of input text in the full context of all tokens before and after, hence the name: Bidirectional Encoder Representations from Transformers (BERT).

BERT models are usually pre-trained on a large corpus of text, then fine-tuned for specific tasks. Consequently, text classification with BERT can be done by fine-tuning a pre-trained BERT by feeding its vector-space representations of text into a classifier trained on domain data [14]. For this reason, the BERT family of models has become a standard approach for the training of task-specific models [15]. Pretraining BERT on large-scale biomedical corpora led to BioBERT, suitable for biomedical text mining tasks [5]. BioBERT is thus available as a pre-trained model that can be transferred to even more specialized models. Similarly, SciBERT is a pretrained model based on BERT and trained using scientific terminology [16]. A recent study that integrated multiple BERT models showed rapid screening, with good accuracy and high sensitivity, of titles and abstracts for SLRs [7]. It was suggested that recent advancements in NLP technology provide a means to rapidly screen literature when updating systematic reviews [7].

A second method explored employs the use of support vector machines (SVM), that is a supervised machine learning algorithm that is commonly used for the purpose of automatic classification of data [17]. In order to use SVMs to classify textual data, such as the titles and abstracts of clinical studies, the texts need to be represented as points in a geometrical space (numerical vectors). This is achieved by decomposing the text of a study into words (tokenization) and word counts with the same linguistic word stem (stemming) that are added up in one entry of the vector representing the study text (context agnostic bag of words representation).

SVMs are based on the idea of dividing a dataset into two classes by determining a linear separation (hyperplane) [17]. The algorithm chooses the hyperplane that results in the greatest distance between the hyperplane and the nearest data points (support vectors) from either training set of the two classes (maximum margin) in order to find the best separation. By constructing more than one SVM and applying advanced analytical methods, data can also be classified into three or more categories simultaneously (multi-label classification). While SVM represent a powerful means for the classification of data, their complexity depends on the size of input data, and they are not optimal for classifying large data sets or text corpora [18]. The use of a multikernel SVM and automatic parameterization markedly improved the speed of training an SVM to manage highly dimensional data with good accuracy [18]. A recent SLR and meta-analysis of publications addressing AI methods used for automated medical literature screening included 86 studies in the review, and 71 in the meta-analysis [19]. This study found that the most commonly used classifier was SVM. Subgroup analyses were performed for different AI algorithms collectively (i.e.; naive Bayes, K-nearest neighbor, perceptron, random forest, convolutional neural networks, radial basis function kernel) and SVM. SVM was considered separately because evidence suggested that SVM classifiers had the best performance for text classification [19]. There were no significant differences in recall, specificity, and precision between SVM and other algorithms collectively [19]. In the present study, fine-tuned BERTs and SVMs were applied to the title and abstract screening phase of two separate clinical SLRs and the results compared with those obtained by humans.

## Methods

Literature searches were performed on EMBASE and run separately for two highly investigated health conditions: metastatic non-small cell lung cancer (mNSCLC) and metastatic castration-resistant prostate cancer (mCRPC). More specifically, the reviews of interest were focused on assessing clinical efficacy and safety of the treatments available for the two indications. For each condition, the title and abstract records from each SLR were reviewed manually, and by two different advanced analytic methods (AAM). A subset of the human labelled records was used to train each of the AAMs. For the mNSCLC study, the reviewers had 2 to 10 years of experience in conducting SLRs. For the mCRPC study, the review was conducted by post graduates in Pharmacy with 2–4 years of experience and quality check was conducted by researchers with 4–5 years of experience. With each approach, including manual review, a classification status (ie; include/exclude) and reason for the exclusion was provided for each record. Inclusion and exclusion status were based on protocol pre-defined study selection criteria: population, intervention, comparator, outcomes and study design (PICOS). The criteria are summarized in Table 1.

**Table 1.**
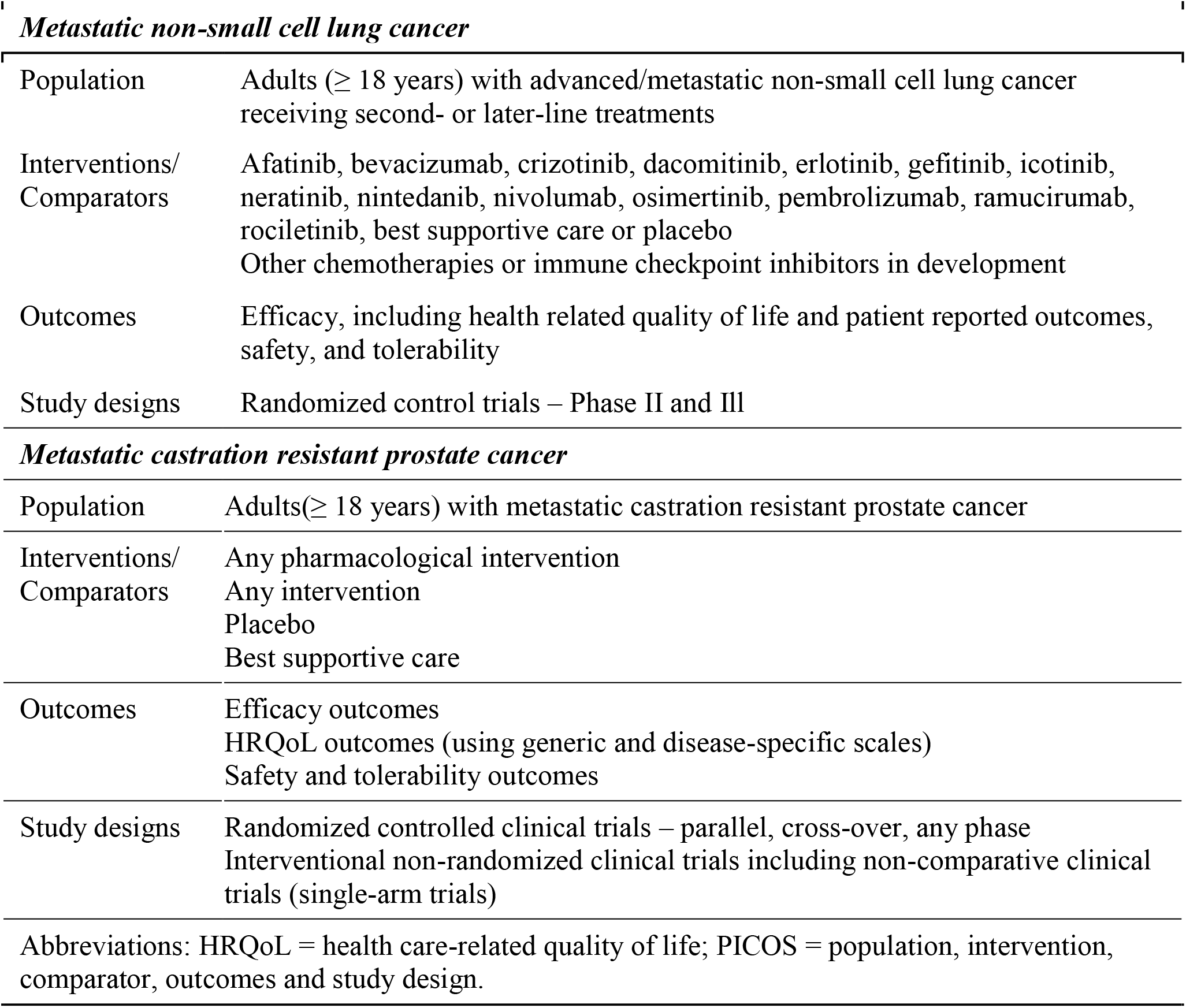
PICOS inclusion criteria

***Metastatic non-small cell lung cancer***
|  |  |
| --- | --- |
| Population | Adults ($\geq 18$ years) with advanced/metastatic non-small cell lung cancer receiving second- or later-line treatments |
| Interventions/<br>Comparators | Afatinib, bevacizumab, crizotinib, dacomitinib, erlotinib, gefitinib, icotinib, neratinib, nintedanib, nivolumab, osimertinib, pembrolizumab, ramucirumab, rociletinib, best supportive care or placebo<br>Other chemotherapies or immune checkpoint inhibitors in development |
| Outcomes | Efficacy, including health related quality of life and patient reported outcomes, safety, and tolerability |
| Study designs | Randomized control trials – Phase II and III |
| ***Metastatic castration resistant prostate cancer*** |  |
| Population | Adults($\geq 18$ years) with metastatic castration resistant prostate cancer |
| Interventions/<br>Comparators | Any pharmacological intervention<br>Any intervention<br>Placebo<br>Best supportive care |
| Outcomes | Efficacy outcomes<br>HRQoL outcomes (using generic and disease-specific scales)<br>Safety and tolerability outcomes |
| Study designs | Randomized controlled clinical trials – parallel, cross-over, any phase<br>Interventional non-randomized clinical trials including non-comparative clinical trials (single-arm trials) |

Abbreviations: HRQoL = health care-related quality of life; PICOS = population, intervention, comparator, outcomes and study design.

AAM-1 was the transfer learning approach. This involved using SciBERT and BioBERT, two publicly available open-source, pretrained models specialized in scientific and biomedical literature. These models were fine-tuned and trained using documents from each of the two literature searches. Learned word representation was extracted and used to train the model for a specific question. The two research topics, mNSCLC and mCRPC, were assessed separately, with a different model created for each subtask of accept, reject and each reason for exclusion. AAM-2 was an AI tool combining SVM with other advanced analytic methods. Between 25% and 30% of the studies from the literature search were used for training, from which more than 10,000 words were tokenized, based on stem forms of the words, and counted into a vector for each study. These were used to determine the hyperplanes to classify each study as “accept” or “reject.” The reasons for exclusion were modeled as multiple categories in a multi-label classification. The algorithm produced a confidence value between 0 and 1 along with the decision. With that, it was possible to abstain from making an automatic decision if the confidence values for all classes were below a certain threshold (i.e. 0.1). This resulted in a few documents which are deemed “unclassifiable”.

### Assessments

Results from the automated reviews were compared to those of the human-conducted classifications. Confusion matrices were created for each of the two SLRs in order to assess how well each of the models performed and to identify the errors made by each model, as shown in Table 2. A confusion matrix summarizes the classification performance of a classifier with respect to the test data and is generally laid out as a table, with each class listed along the columns and row headers. Each cell of the table then contains the numbers of items in each class correctly classified and those erroneously categorized to the other classes [20, 21]. Columns represent the totals of the manual results and rows the totals of the automated results for each class. Therefore, each column total represents the classification for each class as determined by a human reviewer. The results obtained by humans served as the baseline against which the results of the two AAM models were compared.

**Table 2.**
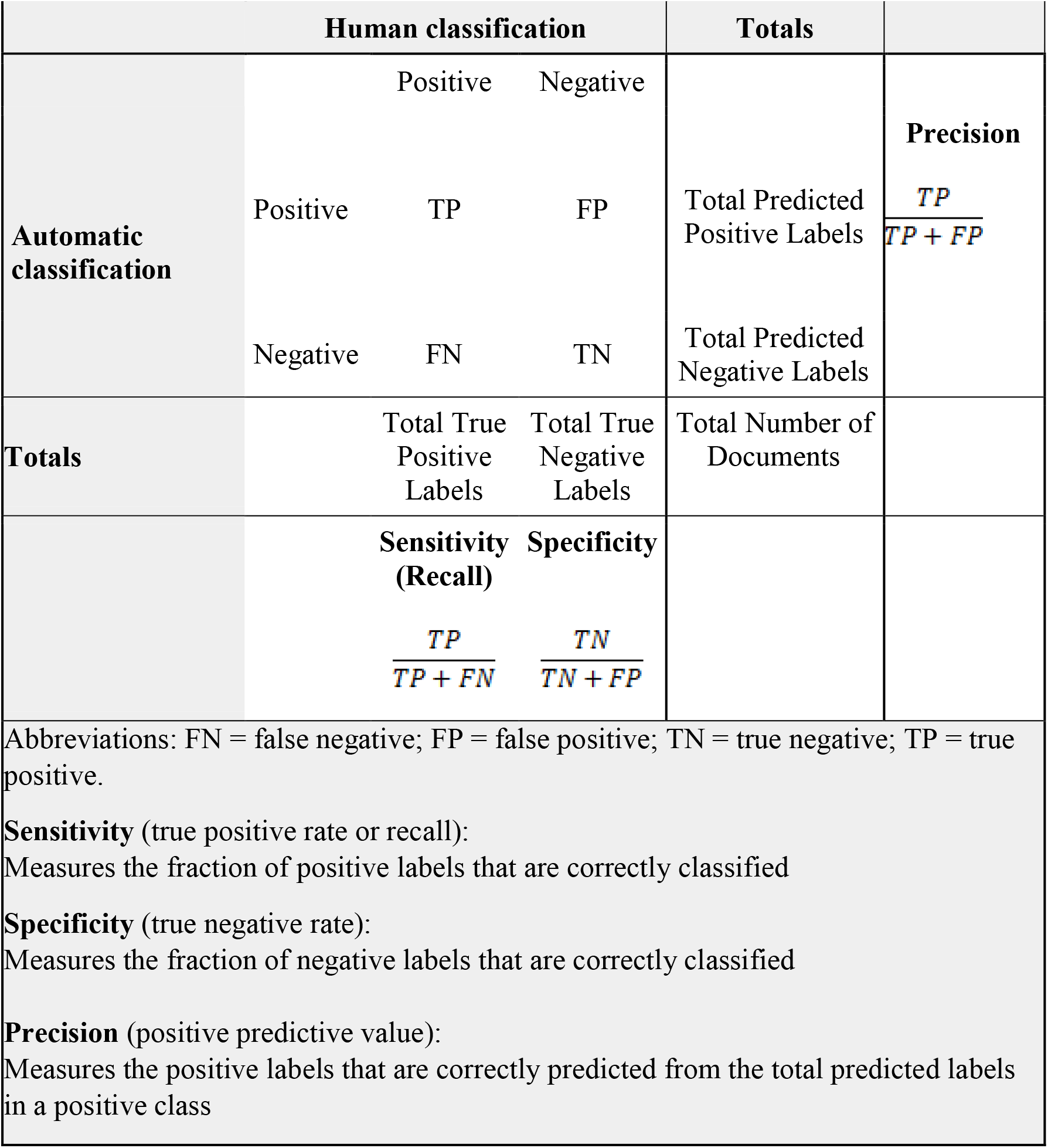
Confusion matrix for binary classification of positive and negative classes

A special case of the confusion matrix is the binary matrix that classifies positive and negative classes to identify true negatives (TN), true positives (PN, false negatives (FN) and false positives (FN), which is depicted in Table 2 [21]. As shown in Table 2, these classifications may be used to calculate performance metrics of the AAMs. Specificity (true negative rate) is a measure of the fraction of negative labels that are correctly classified, sensitivity (true positive rate or recall) is a measure of the fraction of positive labels that are correctly classified, and precision (positive predictive value) is a measure of the positive labels that are correctly predicted from the total predicted labels in a positive class. Multiclass matrices were resolved into a series of binary matrices for each of the accept/reject parameters, in order to calculate a range of performance metrics. Additional metrics, summarized in Table 3, were also used to evaluate the models:

**Table 3.** Assessments

| **Table 3. Assessments** |  |  |  |  |
| --- | --- | --- | --- | --- |
|  | **ROC-AUC** | **F1** | **MCC** | **WSS@95** |
| Definition | Probability that the classifier will rank a randomly chosen positive instance (accept) higher than a randomly chosen negative instance (reject) | Measure of the accuracy of binary (two-class) classification | Measure of the quality of binary (two-class) classification | Work saved at a recall of 95% as the percentage of papers that meet the original search criteria that the reviewers do not have to read |
| Score range | 0 to 1 | 0 to 1 | -1 to 1 | -1 to +1 |
| Working/useful classifier | > 0.8 | Higher is better | Higher is better | >0 |
| Classifier works | > 0.5 | F1 score > F1 score for random classification | > 0 | >0 |
| Random classification | 0.5 | depends on prevalence (inclusion rate):<br>≥ 0 | 0 | 0 |
| Classifier not useful | < 0.5 | F1 score ≤ F1 score for random classification | < 0 | ≤0 |
| Abbreviations: AAM = advanced analytic methods; $F1 = (2 * \text{precision} * \text{recall}) / (\text{precision} + \text{recall})$; MCC = Matthews Correlation Coefficient; measures quality to binary classification; ROC-AUC = receiver-operator characteristic – area under curve; WSS@95 = work saved over sampling at 95% recall. | | | | |

Receiver operating characteristic curve – area under the curve (ROC-AUC) is used to determine how well a model is able to classify items. The ROC curve is a plot of the true positive rate (recall, sensitivity) plotted against the false positive rate, also referred to as (1 − specificity). A ROC-AUC value approaching 1.0 indicates excellent classification skills, whereas one of 0.5 is representative of random guessing, and below that indicates classification that is poorer than guesswork.

The F1 metric is the weighted harmonic mean of a test’s precision and recall. Recall is the fraction of relevant documents that are successfully retrieved, and precision is the fraction of retrieved documents that are relevant to the query. Thus, a result with high precision suggests that the retrieved documents would be highly relevant, whereas a high recall suggests that most, if not all, relevant documents would be retrieved. A high F1 score suggests an acceptable balance between specificity and relevance. To interpret whether a F1 score is “good” or “bad”, we can compare it to a F1 score calculated on the prevalence (positive class: include) of the data (random classification).

The Matthews Correlation Coefficient (MCC) is a measure of the quality of a binary classification. This is a measure of the correlation between observed and predicted classification. A score of 1 indicates perfect correlation, 0 means the prediction is no better than random prediction, and −1 indicates complete disagreement between predicted and observed classifications.

Work Saved over Sampling (WSS) is a metric designed to measure how much future work the reviewers could save for each review. Work saved is defined as the percentage of papers that meet the original search criteria that, because they were screened by the AAM, they do not have to be read by the reviewers [22]. In order for the AAM to provide an advantage, the work saved should be greater than that achieved by random sampling for a given level of recall. The WSS is determined by the formula:

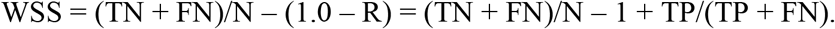

Setting recall at 0.95 results in the formula:

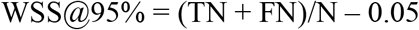

where TP and FN were previously defined above, R is recall, and N is the total number of samples in the test set [22]. Work saved over sampling at 95% recall (WSS@95%) was used as the measure of value to the review process.

In addition, we have performed additional calculations on the time saved with the implementation of the AAMs in the SLR process. The time to complete automated review (TCAR) included the time needed to prepare the training datasets, to train the models and to get the classification for the test set, and is defined by the following formula:

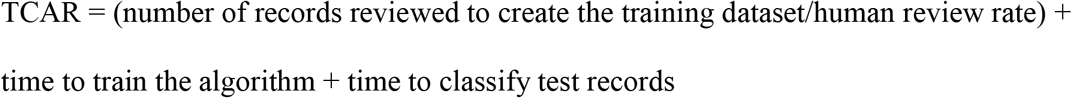

In the formula, human review rate equals the number of records reviewed by humans per time unit. For the analysis presented, we have assumed a human review rate of 40 records per hour.

## Results

### A. Metastatic non-small cell lung cancer

There was a total of 7845 records analyzed during the human-conducted review. Of these, 2025 were used to train both AAM-1 and AAM-2. The remaining 5820 records were analyzed by AAM-1 and AAM-2 using the protocol-predefined criteria. The human-conducted classification took 196 hours to complete, whereas TCAR for AAM-1 and AAM-2 was 52 hours each.

### Binary classification

Human-based classification resulted in acceptance of 8% (440 items) and rejection of 92% (5380 items) of the records. AAM-1 correctly accepted 6% of records, correctly rejected 79% and wrongly classified 15% of the records (Fig 1). As shown in the confusion matrix for binary classification (Table 4), there were 377 records that were correctly accepted and 4626 that were correctly rejected.

**Table 4.** Confusion matrix for binary classification for AAM-1 for mNSCLC

|  |  |  |  |  |  |
| --- | --- | --- | --- | --- | --- |
| **Human classification** |  |  |  |  |  |
| **Automatic classification** |  | Accept | Reject | Total | **Precision** |
|  | Accept | 377 | 754 | 1131 | 0.33 |
|  | Reject | 63 | 4626 | 4689 |  |
|  | Total | 440 | 5380 | 5820 |  |
|  |  | Sensitivity (Recall) | Specificity |  |  |
|  |  | 0.86 | 0.86 |  |  |
| Abbreviations: AAM = advanced analytic methods; mNSCLC = metastatic non-small cell lung cancer. |  |  |  |  |  |

**Table 5.** Confusion matrix for binary classification for AAM-2 for mNSCLC

AAM-2 correctly accepted 6% (361 records) and correctly rejected 82% of the records (4786 record), which means that 12% were wrongly classified (Table 5).
|  |  |  |  |  |  |
| --- | --- | --- | --- | --- | --- |
| **Human classification** |  |  |  |  |  |
| **Automatic classification** |  | Accept | Reject | Total | **Precision** |
|  | Accept | 361 | 594 | 955 | 0.38 |
|  | Reject | 79 | 4786 | 4865 |  |
|  | Total | 440 | 5380 | 5820 |  |
|  |  | Sensitivity (Recall) | Specificity |  |  |
|  |  | 0.82 | 0.89 |  |  |
| Abbreviations: AAM = advanced analytic methods; mNSCLC = metastatic non-small cell lung cancer. |  |  |  |  |  |

[**Fig 1. Disposition of records for the mNSCLC data set**. This schematic drawing shows the proportions of records that were used as a training set, and the proportions rejected, accepted, and classified incorrectly by humans, and by AAM-1 and AAM-2. The times needed to complete the reviews are also shown. Each full square represents 10 hours.]

Based on the data presented in the binary classification confusion matrices giving the TN, FN, FP, and TP values, the assessment metrics, ROC-AUC, F1, MCC and WSS were determined(Table 6). Both AAMs performed similarly. Both methods had ROC-AUC values >0.8, indicating that they both are useful classifiers. The F1 and MCC values indicate that the methods accurately perform binary classifications with a high quality. WSS@95% value was 0.76 for AAM-1 and 0.79 for AAM-2, suggesting a substantial savings in time in performing future classifications.

**Table 6.**
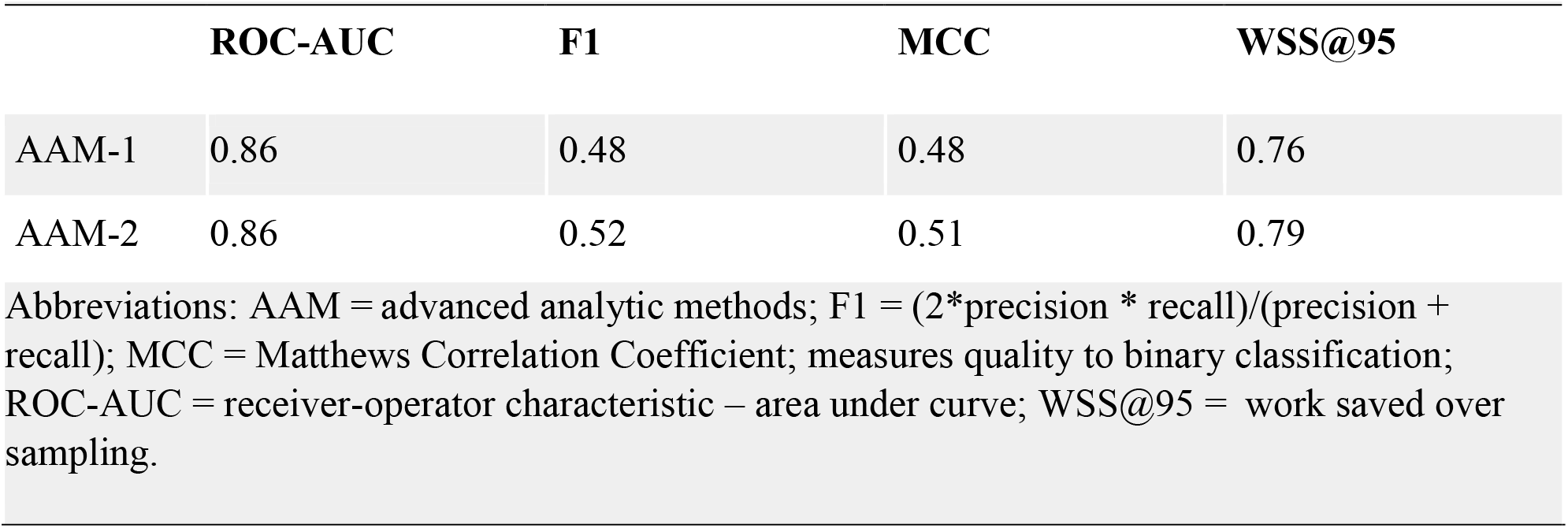
Assessment metrics for binary classification of records for metastatic non-small cell lung cancer

|  | ROC-AUC | F1 | MCC | WSS@95 |
| --- | --- | --- | --- | --- |
| AAM-1 | 0.86 | 0.48 | 0.48 | 0.76 |
| AAM-2 | 0.86 | 0.52 | 0.51 | 0.79 |

Abbreviations: AAM = advanced analytic methods; $F1 = (2 * \text{precision} * \text{recall}) / (\text{precision} + \text{recall})$ ; MCC = Matthews Correlation Coefficient; measures quality to binary classification; ROC-AUC = receiver-operator characteristic – area under curve; WSS@95 = work saved over sampling.

### Multilabel classification

For multilabel classification, the records were also classified into five categories simultaneously, with classes being “accept” and “reject” with reasons for rejection given as “population”, “intervention”, “outcome”, and “study design” with results presented in a multi-classification confusion matrix for AAM-1 (Table 7) and AAM-2 (Table 8). The items that were classified correctly are represented in the shaded cells in a diagonal line in the tables.

**Table 7.**
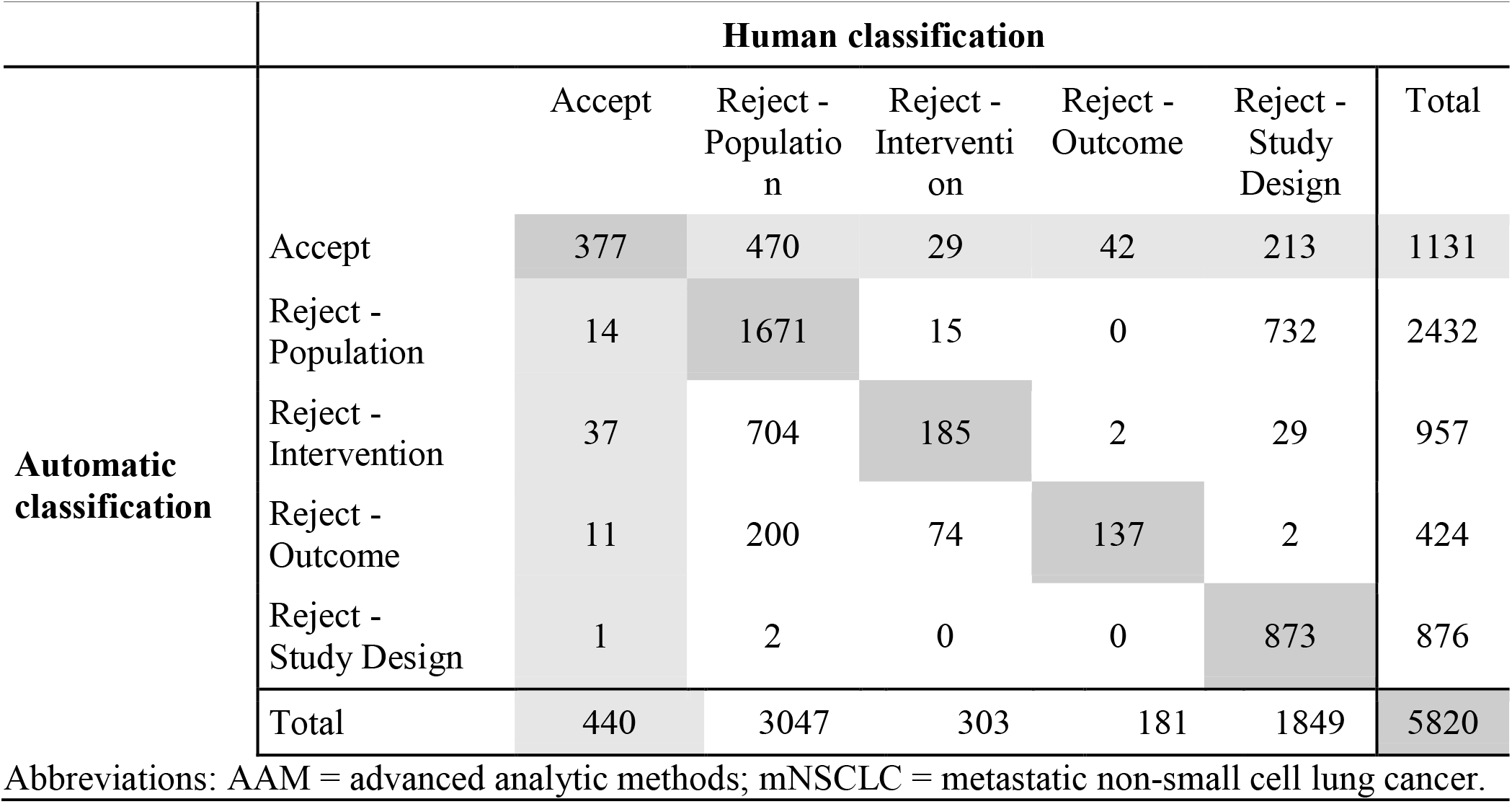
AAM-1 (mNSCLC): Confusion Matrix for multiclass classification: Accept / Reject with Reason for Exclusion.

**Table 8.**
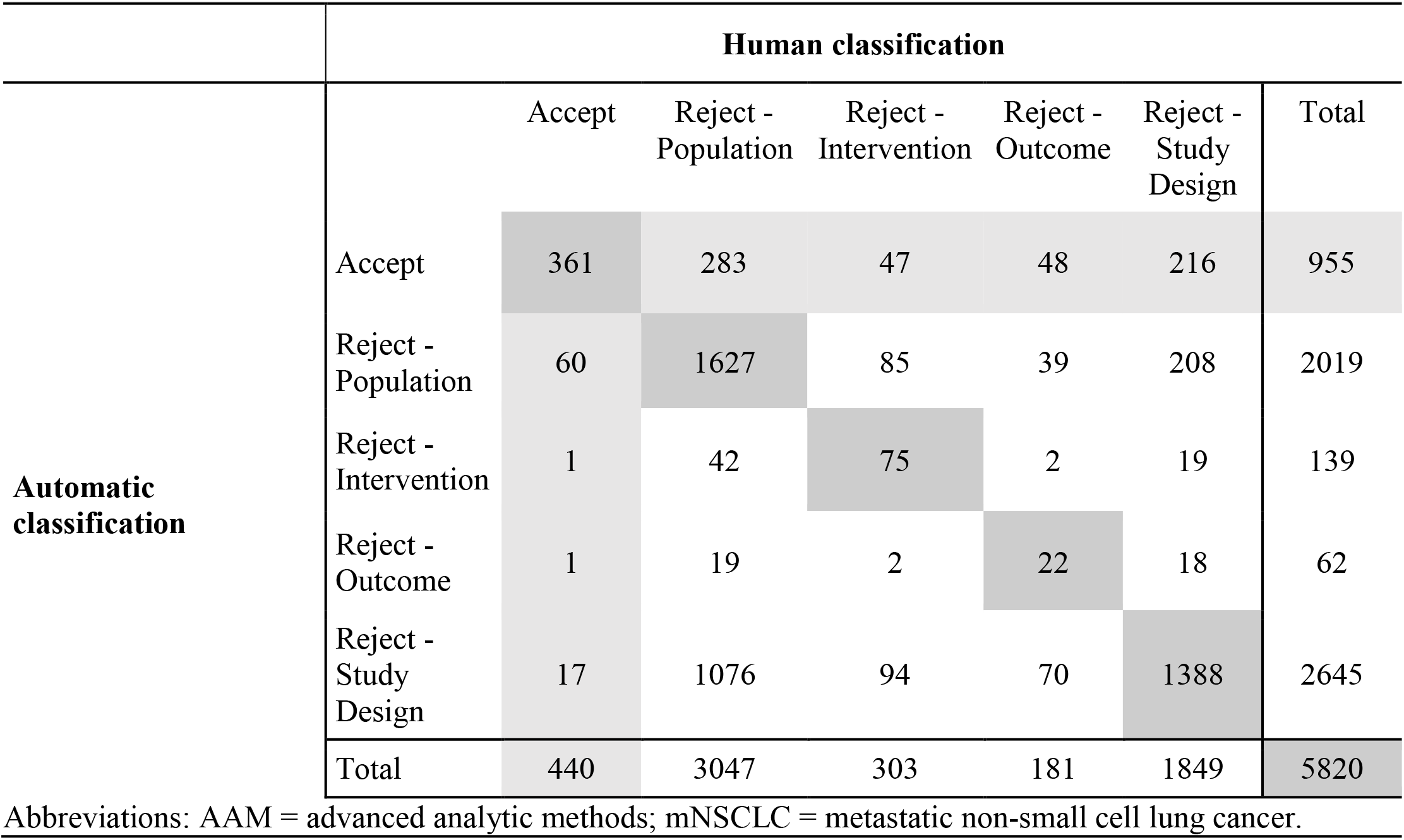
AAM-2 (mNSCLC): Confusion Matrix for multiclass classification: Accept / Reject with Reason for Exclusion.

A total of 2866 records (62%) were rejected for the correct reasons by AAM-1 (Table 7). Among these, 1671 of 3047 were correctly rejected for population, 185 of 303 were correctly rejected for intervention, 137 of 181 were correctly rejected for outcomes, and 873 of 1849 were correctly rejected for study design (Table 7).

Of those correctly rejected by AAM-2, the correct rejection reason was attributed to 3112 (65%) records (Table 8). Among these, 1627 of 3047 were correctly rejected for population, 75 of 303 were correctly rejected for intervention, 22 of 181 were correctly rejected for outcomes, and 1388 of 1849 were correctly rejected for study design (Table 8).

Both methods, AAM-1 and AAM-2, performed similarly. Both methods accept the vast majority of the true accepts (i.e. only few false negatives) while also accepting a substantial amount of true rejects (false positives). Confusion matrices show that approximately 90% of misclassified documents are false positives. The ranges given summarize values calculated for the individual reason for exclusion and highlight that the predictions performance is not consistent across the reasons for exclusion. Both methods had ROC-AUC values ≥0.6, indicating that both classifiers have limited prediction power (Table 9). Thus, predicting the reasons to reject is better than guessing but has limited accuracy with both methods. The performance metrics determined for each of the individual exclusion reasons for the multiclass matrices were determined from binary matrices derived from each of the multiclass matrices (Supplemental Results).

**Table 9.**
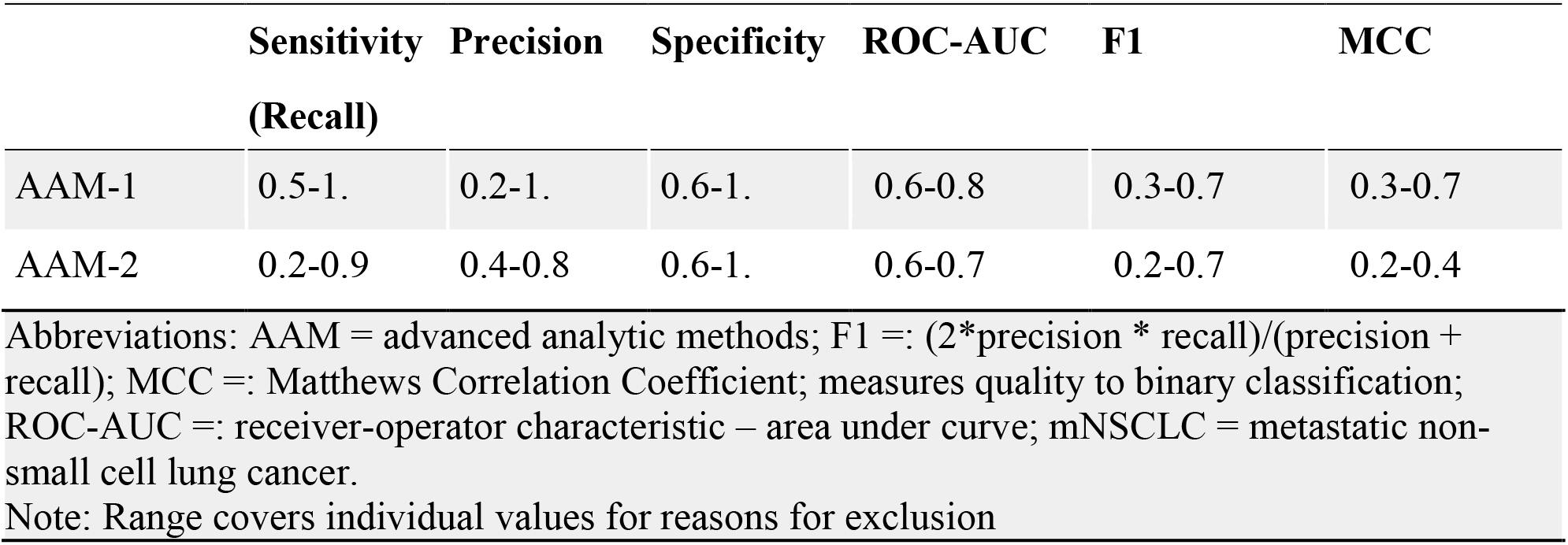
Performance for multi-label classification (mNSCLC)

|  | Sensitivity<br>(Recall) | Precision | Specificity | ROC-AUC | F1 | MCC |
| --- | --- | --- | --- | --- | --- | --- |
| AAM-1 | 0.5-1. | 0.2-1. | 0.6-1. | 0.6-0.8 | 0.3-0.7 | 0.3-0.7 |
| AAM-2 | 0.2-0.9 | 0.4-0.8 | 0.6-1. | 0.6-0.7 | 0.2-0.7 | 0.2-0.4 |

Abbreviations: AAM = advanced analytic methods; F1 =: (2\*precision \* recall)/(precision + recall); MCC =: Matthews Correlation Coefficient; measures quality to binary classification; ROC-AUC =: receiver-operator characteristic – area under curve; mNSCLC = metastatic non-small cell lung cancer. Note: Range covers individual values for reasons for exclusion

## B. Metastatic prostate cancer

There was a total of 3408 records, of which 974 were used to train both AAM-1 and AAM-2, and 2434 were classified by humans and by AAM-1 and AAM-2. The human-conducted classification took 85 hours to complete, whereas TCAR for either AAM-1 or AAM-2 was 25 hours each.

### Binary classification

Human-based classification accepted 26% (638 records) and rejected 74% (1796 records) of the records. AAM-1 correctly accepted 23% (550 records), correctly rejected 62% (1521 records)and wrongly classified 15% (363 records) (Table 10; Fig 2).

[**Fig 2. Disposition of records for the mNSCLC data set**. This schematic drawing shows the proportions of records that were used as a training set, and the proportions rejected, accepted,and classified incorrectly by humans, and by AAM-1 and AAM-2. The times needed to complete the reviews are also shown. Each full square represents 10 hours.]

**Table 10.**
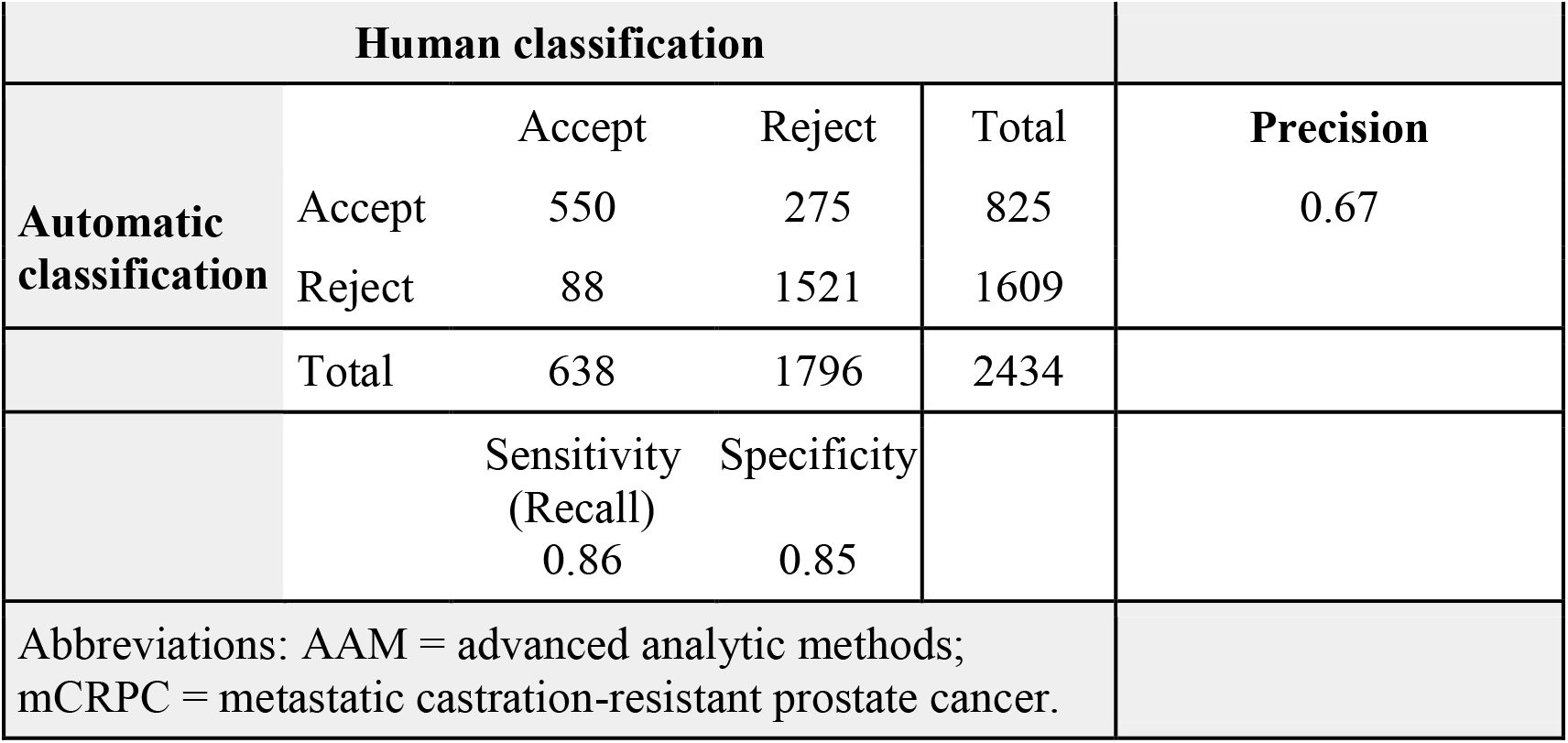
Confusion matrix for binary classification for AAM-1 for mCRPC

AAM-2 correctly accepted 20% (486 records), correctly rejected 66% (1608 records), and wrongly classified 13% (315 records). In addition, there were 25 records (0.6%) that were not classified by AAM-2 (Table 11, Fig 2).

**Table 11.**
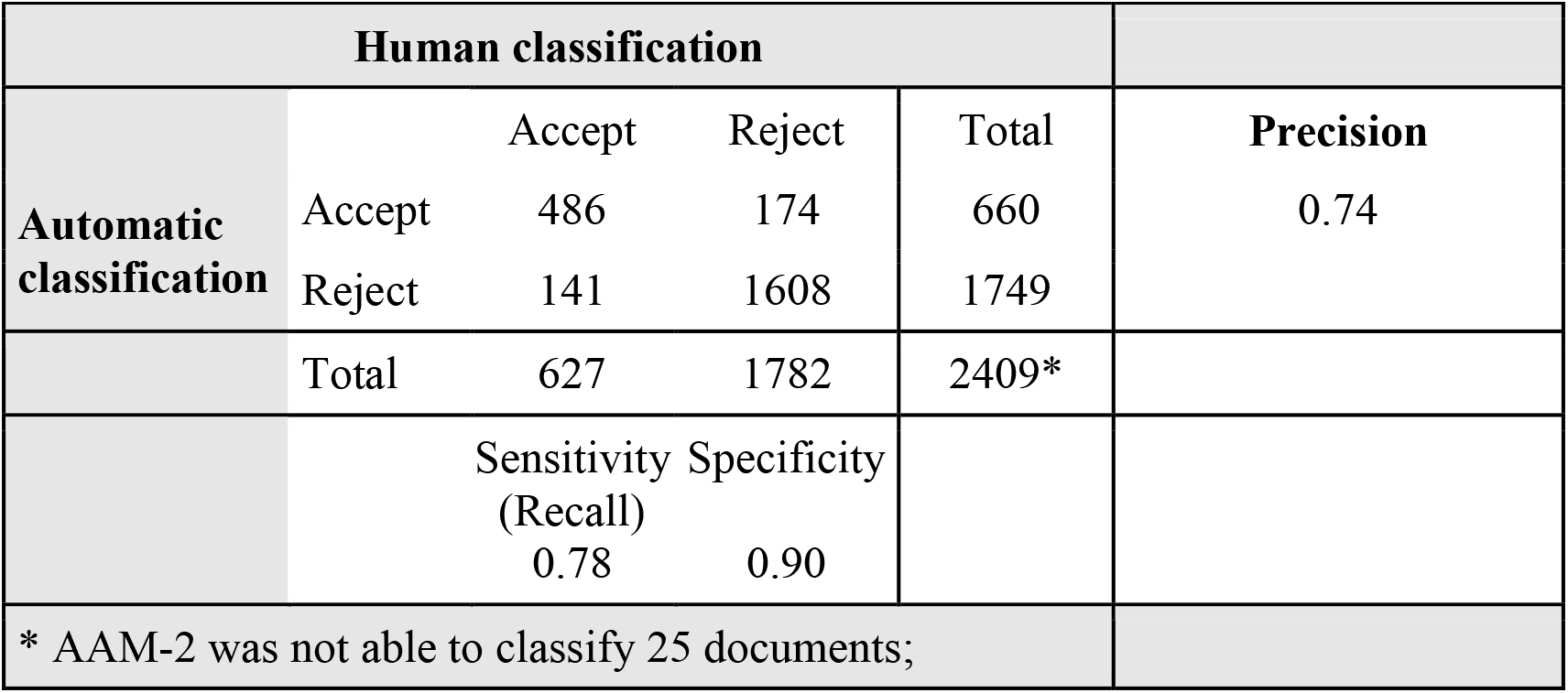
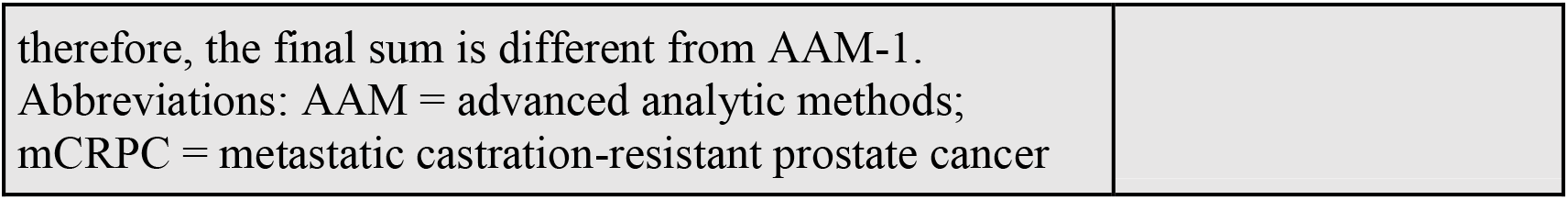
Confusion matrix for binary classification for AAM-2 for mCRPC

The data presented in the binary classification confusion matrices for giving the TN, FN, FP, and TP values, the assessment metrics, ROC-AUC, F1, MCC and WSS, were determined for prostate cancer (Table 12). Both methods, AAM-1 and AAM-2, performed similarly. However, AAM-1 had considerably more false positives than false negatives, whereas AAM-2 had an equivalent number of false positives and false negatives. Both methods had ROC-AUC values >0.8, indicating that they both are useful classifiers. The F1 and MCC values indicate that the methods accurately perform binary classifications with a high quality. AAM-1 and AAM-2 were associated with WSS@95% values of 0.61 for AAM-1 and 0.68 for AAM-2, suggesting a substantial savings in time in performing future classifications.

**Table 12.**
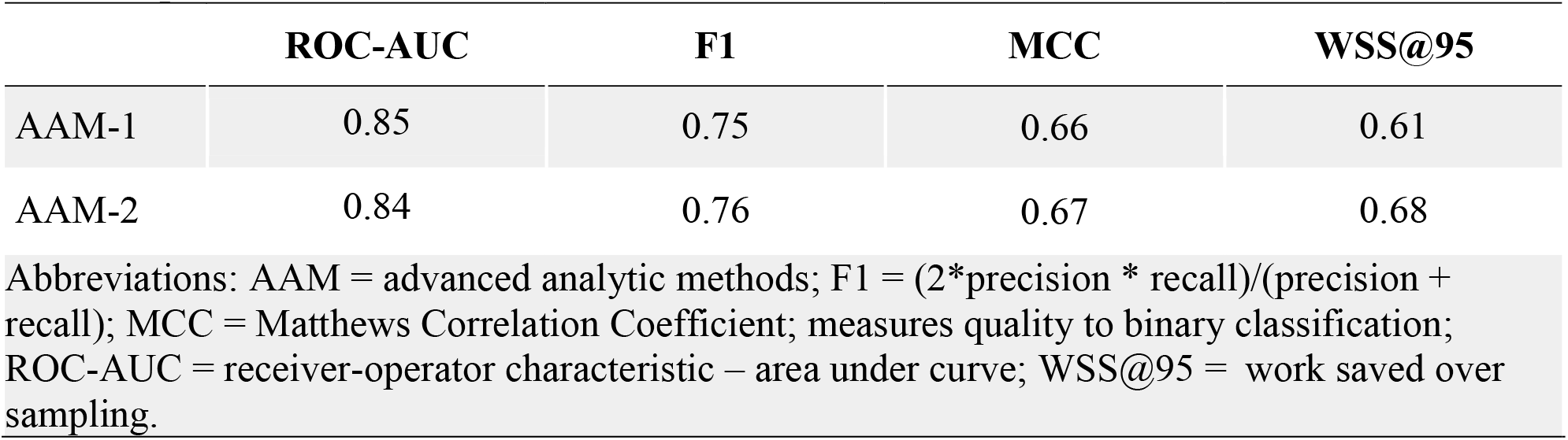
Assessment metrics for binary classification of records for metastatic castration-resistant prostate cancer

|  | ROC-AUC | F1 | MCC | WSS@95 |
| --- | --- | --- | --- | --- |
| AAM-1 | 0.85 | 0.75 | 0.66 | 0.61 |
| AAM-2 | 0.84 | 0.76 | 0.67 | 0.68 |

Abbreviations: AAM = advanced analytic methods; F1 = $(2 * \text{precision} * \text{recall}) / (\text{precision} + \text{recall})$ ; MCC = Matthews Correlation Coefficient; measures quality to binary classification; ROC-AUC = receiver-operator characteristic – area under curve; WSS@95 = work saved over sampling.

### Multilabel classification

The records were also classified into four categories simultaneously, with classes being “accept” and “reject” with reasons for rejection given as “population”, “intervention”, “study design” Results are presented in a multi-classification confusion matrix for AAM-1 (Table 13) and AAM-2 (Table 14). Although “outcomes” were part of eligibility criteria defined in the protocol for this condition, this class was not used as an exclusion reason in the title and abstract review level. For records rejected by AAM-1, 12 of 202 were correctly rejected for population, 0 of 5 were correctly rejected for intervention /comparator, and 1363 of 1589 were correctly rejected for study design (Table 13). For records rejected by AAM-2, 25 of 196 were correctly rejected for population, 0 of 5 were correctly rejected for intervention /comparator, and 1454 of 1581 were correctly rejected for study design (Table 14).

**Table 13.**
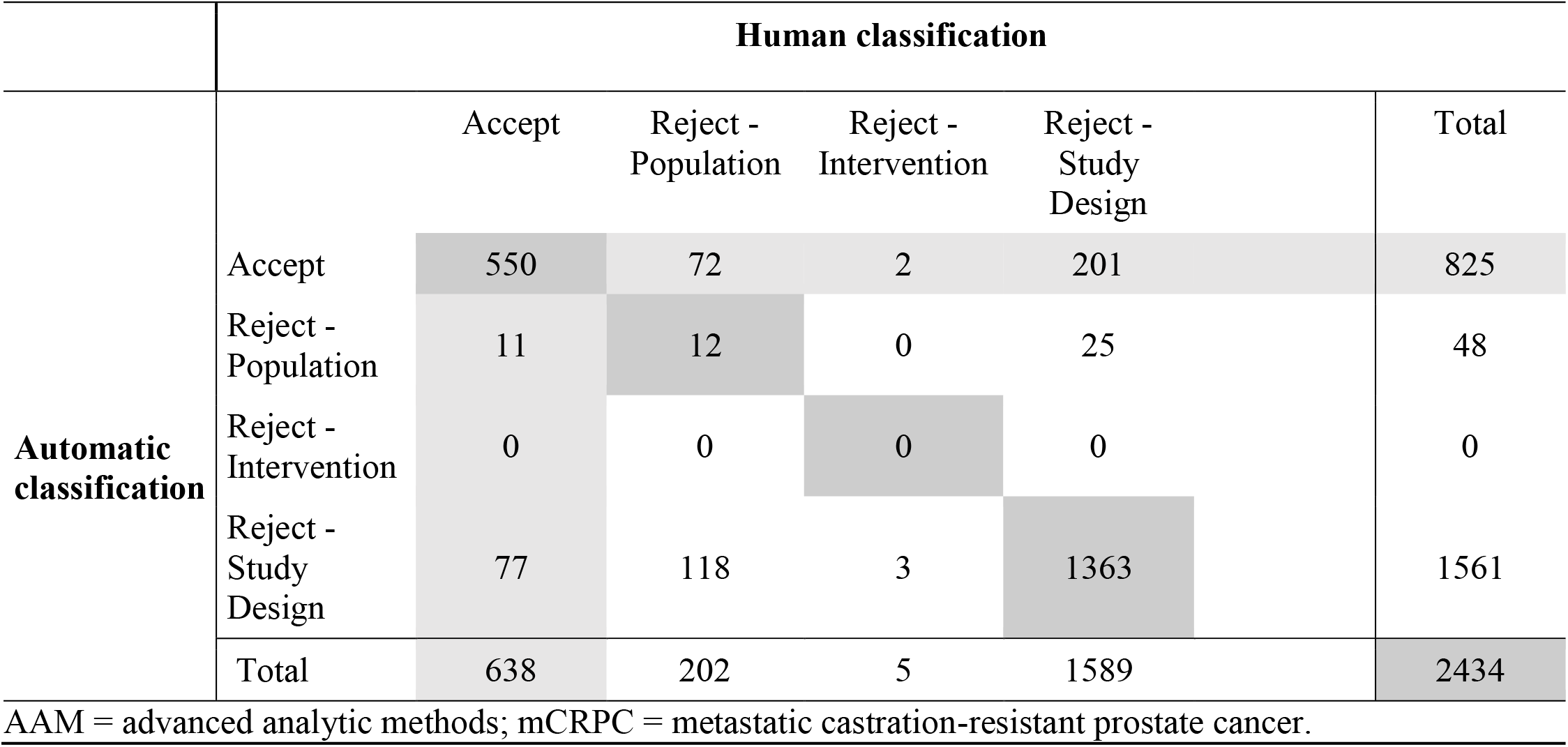
Confusion Matrix for multiclass classification for mCRPC: AAM-1: Accept / Reject with Reason for Exclusion.

**Table 14.**
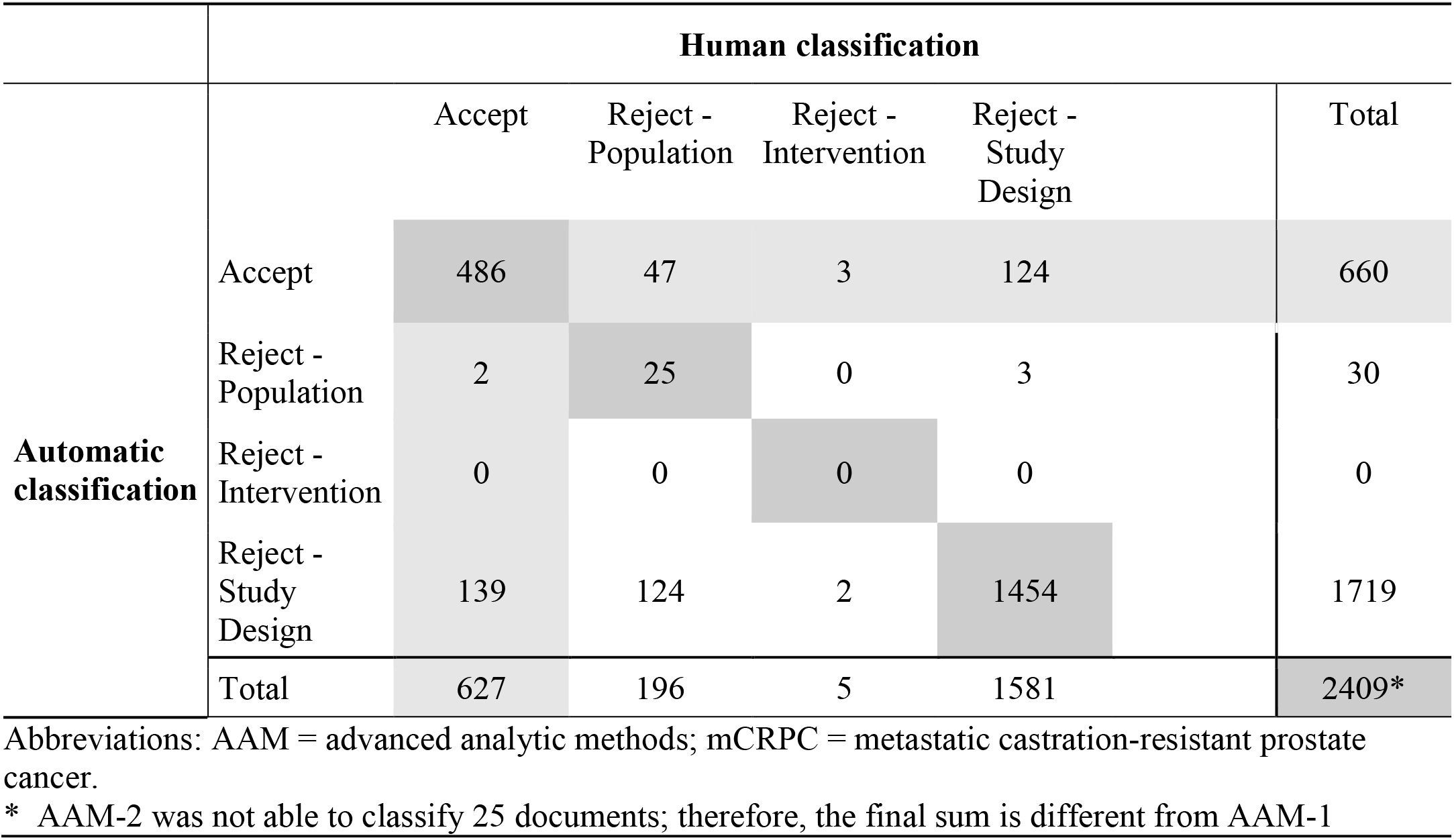
Confusion Matrix for multiclass classification for mCRPC: AAM-2: Accept / Reject with Reason for Exclusion.

Both methods, AAM-1 and AAM-2, performed similarly. The ranges given summarize values calculated for the individual reason for exclusion and highlight that the predictions performance is not consistent across the reasons for exclusion. Both methods had ROC-AUC values >0.6 for multilabel classification, indicating that both AAMs have limited prediction power. Thus,predicting the reasons to reject is better than guessing but has limited accuracy with both methods. Moreover, for both AAM-1 and AAM-2, the classifier for exclusion reason Intervention did not work very well, which resulted in the low values (ie; 0) at the start of each range for the metrics shown in Table 15. The performance metrics determined for each of the individual exclusion reasons for the multiclass matrices were determined from binary matrices derived from each of the multiclass matrices (Supplemental Results).

**Table 15.**
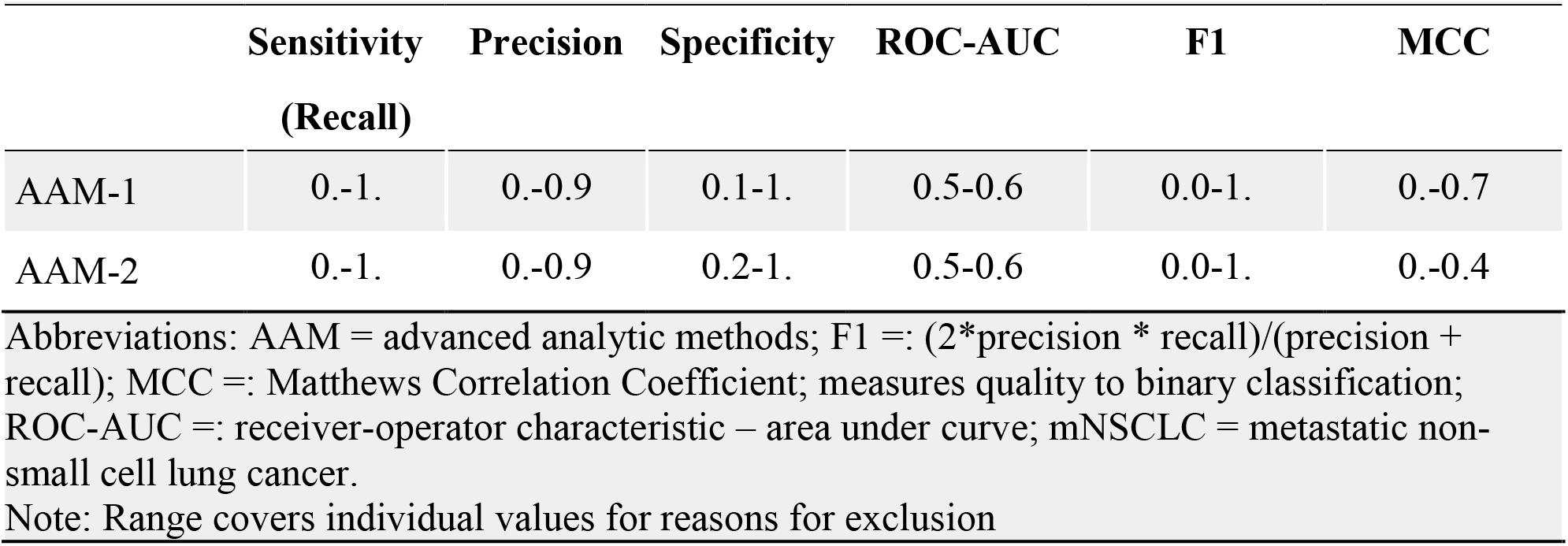
Performance for multi-label classification (mCRPC)

|  | **Sensitivity**<br>**(Recall)** | **Precision** | **Specificity** | **ROC-AUC** | **F1** | **MCC** |
| --- | --- | --- | --- | --- | --- | --- |
| AAM-1 | 0.-1. | 0.-0.9 | 0.1-1. | 0.5-0.6 | 0.0-1. | 0.-0.7 |
| AAM-2 | 0.-1. | 0.-0.9 | 0.2-1. | 0.5-0.6 | 0.0-1. | 0.-0.4 |

Abbreviations: AAM = advanced analytic methods; F1 =: $(2 * \text{precision} * \text{recall}) / (\text{precision} + \text{recall})$ ; MCC =: Matthews Correlation Coefficient; measures quality to binary classification; ROC-AUC =: receiver-operator characteristic – area under curve; mNSCLC = metastatic non-small cell lung cancer.
Note: Range covers individual values for reasons for exclusion

## Discussion

We assessed how two advanced analytic methods performed in comparison to humans in the review of titles and abstracts during the SLR process for two health conditions. The automated reviews presented herein were performed in order to assess the speed and accuracy of these AAMs in classifying records compared to human reviewers. In general, the machine-learning methods produced results comparable to human review of title and abstracts in considerably less time. The time needed for classification by AAM-1 and AAM-2 was the same within each SLR, and up to four times faster than human reviewers. This represents a substantial saving in time spent, as manual review of abstracts can be a very time-consuming task, taking days to complete. The WSS@95% values for AAM-1 and AAM-2 for both searches indicated predicted work saved of >50%, which is considered to be considerable [22]. Similar results were obtained in the two disease categories, although it might be argued that, based on the F1 scores, that the mNSCLC classifications were not as robust as the mCRPC classifications. In addition, both methods performed well in the binary classification of records, but not as much for providing the reasons for exclusion using the pre-specified PICOS criteria. In a recent meta-analyses of AI algorithms for automated searches, the combined recall, specificity, and precision values were 0.928 [95% confidence interval (CI), 0.878– 0.958], 0.647 (95% CI, 0.442–0.809), and 0.200(95% CI, 0.135–0.287) when achieving maximized recall, and were 0.708 (95% CI, 0.570–0.816), 0.921 (95% CI, 0.824–0.967), and 0.461 (95% CI, 0.375–0.549) when achieving maximized precision [19]. The approach used in the present study was not designed to optimize either recall or precision; however, the values of recall (0.78 to 0.86), specificity (0.85 to 0.90), and precision (0.33 to 0.74) obtained with the binary classification of records in the present study are within the range of values reported by Feng et al. [19].

Although this analysis focuses on clinical RCT evidence in oncology health conditions, the results look promising. We do not know how well the methods would have performed for SLRs focused on other topics, such as economic, epidemiologic, and humanistic, or other health conditions. In the present investigation, although the two AAMs employed behaved similarly within a therapeutic area, there were differences in robustness in the analyses performed for mNSCLC and mCRPC. The reasons for the difference in robustness between the two therapeutic areas are unclear, but may be in part by factors such as size of the search results, the terminologies used, the large imbalance in prevalence of the Accept and Reject classes, or other factors that require further investigation. For example, the imbalance in prevalence of the Accept and Reject classes is much higher for mNSCLC (8% Accepts vs. 92% Rejects) compared to mCRPC (26% Accepts vs. 74% Rejects) indicating that the classifiers do better predictions when imbalances between the classes are smaller. As such, future research should aim to understand how consistent the performance of these AAMs is for reviewing economic, epidemiological, humanistic, treatment patterns and burden of illness evidence in other disease areas.

It should be noted that between 12% and 15% of records were wrongly classified, which is similar to the 10% error rate reported with human reviewers [23]. As most of these records were false positives, they were not excluded, but would have been processed in the next phase; therefore, they do not represent information that was lost. With the false negatives, there would be no next steps, and we would not recover that information. Overall, it appears that results obtained with AAMs may approach the accuracy provided by human reviewers, but there remains room for improvement.

Limitations of the study include the fact that human-run literature searches are considered a “gold standard”, but humans also make mistakes in classification [23], suggesting that there is no one single source of a true result. Consequently, machines may be taught with errors and/or comparisons with machine results may also contain errors. Further investigation is needed to understand what would be the result from a conflict resolution between the automatic classification and the initial human classification. Also, while records could have been excluded as having irrelevant outcomes during the human review of mCRPC records due to lack of reporting of outcomes pre-specified in the SLR protocol, this exclusion reason was not applied at the title and abstract screening level. This practice could sometimes be used by literature review teams that are careful about wrongfully excluding records that meet other inclusion criteria but could potentially provide data on outcomes during the next level review of full-text papers.

Consequently, no records classified as having irrelevant outcomes were used to train either of the AAMs that assessed the mCRPC evidence. Nevertheless, the AAMs were still able to produce results comparable to the human reviewers.

Another potential limitation is that the SVM found 25 mCRPC records to be very close to the decision boundary (hyperplane), so automatic classification was non-conclusive for records with a confidence value below 0.1. These examples should be reviewed manually. The AAMs appeared to work better for accept/reject classification than for assigning reasons for the rejected records. In addition, automated methods require training data. The impact of size, nature and generation process of the training set, as well as the impact of the overall size of the database, is not entirely clear.

Lastly, although, the WSS@95% is a largely documented metric in the literature for estimating resource savings related to literature reviews, it focuses only on the effort saved when comparing the work done by a machine versus what a human would have done. It does not take into account the time needed to prepare the training datasets or train the AAMs, which could be significant depending on how training data is required. For that reason, we propose an additional metric that captures all the effort expended in completing an automatic review – the time to complete automated review (TCAR). This metric is particularly relevant in cases where training is involved. Our analysis found that a key driver of this metric is how long it takes to create and prepare the training dataset. Consequently, we acknowledge that the human review rate in our work may be conservative for some teams that are experts in literature reviewing. In the cases explored in our research, retrospective data from previously completed SLRs were used, and as such the final status of each record was known prior to preparing the training dataset. In a *de novo* review, such information is not available. As a result, to prepare the training dataset, it may be necessary to screen more records than the ones actually used to train the model.

## Conclusion

Use of advanced analytical methods have the potential to markedly reduce the time required for searching and triaging records during a systematic review. These methods should be tested with literature from other therapeutic areas to evaluate if similar levels of accuracy are found. The use of advanced analytic techniques for the classification of records to support an SLR is promising and should improve with additional techniques and refinements.

## Supporting information

Suppplemental Materials

## Acknowledgments

Evidera PPD was contracted for writing and editorial services. Writing support was provided by Michael H. Ossipov, PhD of Evidera PPD.

