## Supplementary material for "Are Machines-learning Methods More Efficient than Humans in Triaging Literature for Systematic Reviews?": Suppplemental Materials

### Supplemental Results

The multiclass confusion matrices were resolved into a series of binary matrices in order to calculate precision, sensitivity (recall), and specificity for each of the reasons for selection or rejection. Thus, for the confusion matrix presented in Table 7, reproduced herein, for AAM-1 (mNSCLC), new binary confusion matrices for the individual exclusion reasons (ER) of “Population” (Supplemental Tables 1), “Intervention” (Supplemental Table 2), “Outcome” (Supplemental Table 3), and “Study Design” (Supplemental Table 4) were constructed, using the parameters outlined in red in Table 7. The element in the “Accept” row and “Accept” column was omitted (i.e.; corresponding to the “True Positive” in Table 4) as it did not represent an ER value. The elements in the “Accept” row across all “Reject” columns were omitted as these contained no predicted ER, only observed ER values (i.e.; corresponding to the “False Positive” in Table 4). The elements in the “Accept” column across all “Reject” were omitted since they contained only predicted ER values, no observed ER values (i.e.; corresponding to the “False Negative” in Table 4). Only those elements that had predicted and observed ER values were used in constructing the binary matrices.

**Table 7. AAM-1 (mNSCLC): Confusion Matrix for multiclass classification: Accept / Reject with Reason for Exclusion.**

| | | Human classification | | | | | |
|---|---|---|---|---|---|---|---|
| | | Accept | Reject - Population | Reject - Intervention | Reject - Outcome | Reject - Study Design | Total |
| **Automatic classification** | Accept | 377 | 470 | 29 | 42 | 213 | 1131 |
| | Reject - Population | 14 | 1671 | 15 | 0 | 732 | 2432 |
| | Reject - Intervention | 37 | 704 | 185 | 2 | 29 | 957 |
| | Reject - Outcome | 11 | 200 | 74 | 137 | 2 | 424 |
| | Reject - Study Design | 1 | 2 | 0 | 0 | 873 | 876 |
| | Total | 440 | 3047 | 303 | 181 | 1849 | 5820 |

Abbreviations: AAM = advanced analytic methods; mNSCLC = metastatic non-small cell lung cancer.

| **Supplemental Table 1. AAM-1 (mNSCLC): Confusion matrix for binary classification: Accept / Reject with Reason for Exclusion: Population** | | | | | |
|---|---|---|---|---|---|
| | | **Human classification** | | | |
| | | Accept | Reject | Total | **Precision** |
| **Automatic classification** | Accept | 1671 | 747 | 2418 | 0.69 |
| | Reject | 906 | 1302 | 2208 | |
| | Total | 2577 | 2049 | 4626 | |
| | | **Sensitivity** 0.65 | **Specificity** 0.64 | | |

Abbreviations: AAM = advanced analytic methods; mNSCLC = metastatic non-small cell lung cancer.

| **Supplemental Table 2. AAM-1 (mNSCLC): Confusion matrix for binary classification: Accept / Reject with Reason for Exclusion: Intervention** | | | | | |
|---|---|---|---|---|---|
| | | **Human classification** | | | |
| | | Accept | Reject | Total | **Precision** |
| **Automatic classification** | Accept | 185 | 735 | 920 | 0.2 |
| | Reject | 89 | 3617 | 3706 | |
| | Total | 274 | 4362 | 4626 | |
| | | **Sensitivity** 0.68 | **Specificity** 0.83 | | |

Abbreviations: AAM = advanced analytic methods; mNSCLC = metastatic non-small cell lung cancer.

| **Supplemental Table 3. AAM-1 (mNSCLC): Confusion matrix for binary classification: Accept / Reject with Reason for Exclusion: Outcome** | | | | | |
|---|---|---|---|---|---|
| | | **Human classification** | | | |
| **Automatic classification** | | Accept | Reject | Total | **Precision** |
| | Accept | 137 | 276 | 413 | 0.33 |
| | Reject | 2 | 4211 | 4213 | |
| | Total | 139 | 4487 | 4626 | |
| | | **Sensitivity** 0.99 | **Specificity** 0.94 | | |
| Abbreviations: AAM = advanced analytic methods; mNSCLC = metastatic non-small cell lung cancer. | | | | | |

| **Supplemental Table 4. AAM-1 (mNSCLC): Confusion matrix for binary classification: Accept / Reject with Reason for Exclusion: Study Design** | | | | | |
|---|---|---|---|---|---|
| | | **Human classification** | | | |
| **Automatic classification** | | Accept | Reject | Total | **Precision** |
| | Accept | 873 | 2 | 875 | 1. |
| | Reject | 763 | 2988 | 3751 | |
| | Total | 1636 | 2990 | 4626 | |
| | | **Sensitivity** 0.53 | **Specificity** 1. | | |
| Abbreviations: AAM = advanced analytic methods; mNSCLC = metastatic non-small cell lung cancer. | | | | | |

For the confusion matrix presented in Table 8, reproduced herein, for AAM-2 (mNSCLC), new binary confusion matrices for the individual exclusion reasons (ER) of "Population" (Supplemental Tables 5), "Intervention" (Supplemental Table 6), "Outcome" (Supplemental Table 7), and "Study Design" (Supplemental Table 8) were constructed, using the parameters outlined in red in Table 8. The element in the "Accept" row and "Accept" column was omitted (i.e.; corresponding to the "True Positive" in Table 5) as it did not represent an ER value. The elements in the "Accept" row across all "Reject" columns were omitted as these contained no predicted ER, only observed ER values (i.e.; corresponding to the "False Positive" in Table 5). The elements in the "Accept" column across all "Reject" were omitted since they contained only predicted ER values, no observed ER values (i.e.; corresponding to the "False Negative" in Table 5). Only those elements that had predicted and observed ER values were used in constructing the binary matrices.

**Table 8. AAM-2 (mNSCLC): Confusion Matrix for multiclass classification: Accept / Reject with Reason for Exclusion.**

| | | **Human classification** | | | | | |
|---|---|---|---|---|---|---|---|
| | | Accept | Reject - Population | Reject - Intervention | Reject - Outcome | Reject - Study Design | Total |
| **Automatic classification** | Accept | 361 | 283 | 47 | 48 | 216 | 955 |
| | Reject - Population | 60 | 1627 | 85 | 39 | 208 | 2019 |
| | Reject - Intervention | 1 | 42 | 75 | 2 | 19 | 139 |
| | Reject - Outcome | 1 | 19 | 2 | 22 | 18 | 62 |
| | Reject - Study Design | 17 | 1076 | 94 | 70 | 1388 | 2645 |
| | Total | 440 | 3047 | 303 | 181 | 1849 | 5820 |

Abbreviations: AAM = advanced analytic methods; mNSCLC = metastatic non-small cell lung cancer.

| **Supplemental Table 5. AAM-2 (mNSCLC): Confusion matrix for binary classification: Accept / Reject with Reason for Exclusion: Population** | | | | | |
|---|---|---|---|---|---|
| **Human classification** | | | | | |
| **Automatic classification** | | Accept | Reject | Total | **Precision** |
| | Accept | 1627 | 332 | 1959 | 0.83 |
| | Reject | 1137 | 1690 | 2827 | |
| | Total | 2764 | 2022 | 4786 | |
| | | **Sensitivity** 0.59 | **Specificity** 0.84 | | |
| Abbreviations: AAM = advanced analytic methods; mNSCLC = metastatic non-small cell lung cancer. | | | | | |

| **Supplemental Table 6. AAM-2 (mNSCLC): Confusion matrix for binary classification: Accept / Reject with Reason for Exclusion: Intervention** | | | | | |
|---|---|---|---|---|---|
| **Human classification** | | | | | |
| **Automatic classification** | | Accept | Reject | Total | **Precision** |
| | Accept | 75 | 63 | 138 | 0.54 |
| | Reject | 181 | 4467 | 4648 | |
| | Total | 256 | 4530 | 4786 | |
| | | **Sensitivity** 0.29 | **Specificity** 0.99 | | |
| Abbreviations: AAM = advanced analytic methods; mNSCLC = metastatic non-small cell lung cancer. | | | | | |

| Supplemental Table 7. AAM-2 (mNSCLC): Confusion matrix for binary classification: Accept / Reject with Reason for Exclusion: Outcome | | | | | |
|---|---|---|---|---|---|
| | | **Human classification** | | | |
| **Automatic classification** | | Accept | Reject | Total | **Precision** |
| | Accept | 22 | 39 | 61 | 0.36 |
| | Reject | 111 | 4614 | 4725 | |
| | Total | 133 | 4653 | 4786 | |
| | | **Sensitivity** 0.17 | **Specificity** 0.99 | | |
| Abbreviations: AAM = advanced analytic methods; mNSCLC = metastatic non-small cell lung cancer. | | | | | |

| Supplemental Table 8. AAM-2 (mNSCLC): Confusion matrix for binary classification: Accept / Reject with Reason for Exclusion: Study Design | | | | | |
|---|---|---|---|---|---|
| | | **Human classification** | | | |
| **Automatic classification** | | Accept | Reject | Total | **Precision** |
| | Accept | 1388 | 1240 | 2628 | 0.53 |
| | Reject | 245 | 1913 | 2158 | |
| | Total | 1633 | 3153 | 4786 | |
| | | **Sensitivity** 0.85 | **Specificity** 0.61 | | |
| Abbreviations: AAM = advanced analytic methods; mNSCLC = metastatic non-small cell lung cancer. | | | | | |

For the confusion matrix presented in Table 13, reproduced herein, for AAM-1 (mCRPC), new binary confusion matrices for the individual exclusion reasons (ER) of "Population" (Supplemental Tables 9), "Intervention" (Supplemental Table 10), and "Study Design" (Supplemental Table 11) were constructed, using the parameters outlined in red in Table 13. The element in the "Accept" row and "Accept" column was omitted (i.e.; corresponding to the "True Positive" in Table 10) as it did not represent an ER value. The elements in the "Accept" row across all "Reject" columns were omitted as these contained no predicted ER, only observed ER values (i.e.; corresponding to the "False Positive" in Table 10). The elements in the "Accept" column across all "Reject" were omitted since they contained only predicted ER values, no observed ER values (i.e.; corresponding to the "False Negative" in Table 10). Only those elements that had predicted and observed ER values were used in constructing the binary matrices.

**Table 13. Confusion Matrix for multiclass classification for mCRPC: AAM-1: Accept / Reject with Reason for Exclusion.**

| | | **Human classification** | | | | |
|---|---|---|---|---|---|---|
| | | Accept | Reject - Population | Reject - Intervention | Reject - Study Design | Total |
| **Automatic classification** | Accept | 550 | 72 | 2 | 201 | 825 |
| | Reject - Population | 11 | 12 | 0 | 25 | 48 |
| | Reject - Intervention | 0 | 0 | 0 | 0 | 0 |
| | Reject - Study Design | 77 | 118 | 3 | 1363 | 1561 |
| | Total | 638 | 202 | 5 | 1589 | 2434 |

AAM = advanced analytic methods; mCRPC = metastatic castration-resistant prostate cancer.

| **Supplemental Table 9. AAM-1 (mCRPC): Confusion matrix for binary classification: Accept / Reject with Reason for Exclusion: Population** | | | | | |
|---|---|---|---|---|---|
| | | **Human classification** | | | |
| | | Accept | Reject | Total | **Precision** |
| **Automatic classification** | Accept | 12 | 25 | 37 | 0.32 |
| | Reject | 118 | 1366 | 1484 | |
| | Total | 130 | 1391 | 1521 | |
| | | **Sensitivity** 0.09 | **Specificity** 0.98 | | |
| Abbreviations: AAM = advanced analytic methods; mNSCLC = metastatic non-small cell lung cancer. | | | | | |

| **Supplemental Table 10. AAM-1 (mCRPC): Confusion matrix for binary classification: Accept / Reject with Reason for Exclusion: Intervention** | | | | | |
|---|---|---|---|---|---|
| | | **Human classification** | | | |
| | | Accept | Reject | Total | **Precision** |
| **Automatic classification** | Accept | 0 | 0 | 0 | n/a |
| | Reject | 3 | 1518 | 1521 | |
| | Total | 3 | 1518 | 1521 | |
| | | **Sensitivity** 0. | **Specificity** 1. | | |
| Abbreviations: AAM = advanced analytic methods; mNSCLC = metastatic non-small cell lung cancer. | | | | | |

| Supplemental Table 11. AAM-1 (mCRPC): Confusion matrix for binary classification: Accept / Reject with Reason for Exclusion: Study Design | | | | | |
|---|---|---|---|---|---|
| **Human classification** | | | | | |
| **Automatic classification** | | Accept | Reject | Total | **Precision** |
| | Accept | 1363 | 121 | 1484 | 0.92 |
| | Reject | 25 | 12 | 37 | |
| | Total | 1388 | 133 | 1521 | |
| | | **Sensitivity** 0.98 | **Specificity** 0.09 | | |
| Abbreviations: AAM = advanced analytic methods; mNSCLC = metastatic non-small cell lung cancer. | | | | | |

For the confusion matrix presented in Table 14, reproduced herein, for AAM-2 (mCRPC), new binary confusion matrices for the individual exclusion reasons (ER) of "Population" (Supplemental Tables 12), "Intervention" (Supplemental Table 13), and "Study Design" (Supplemental Table 14) were constructed, using the parameters outlined in red in Table 13. The element in the "Accept" row and "Accept" column was omitted (i.e.; corresponding to the "True Positive" in Table 11) as it did not represent an ER value. The elements in the "Accept" row across all "Reject" columns were omitted as these contained no predicted ER, only observed ER values (i.e.; corresponding to the "False Positive" in Table 11). The elements in the "Accept" column across all "Reject" were omitted since they contained only predicted ER values, no observed ER values (i.e.; corresponding to the "False Negative" in Table 11). Only those elements that had predicted and observed ER values were used in constructing the binary matrices.

**Table 14. Confusion Matrix for multiclass classification for mCRPC: AAM-2: Accept / Reject with Reason for Exclusion.**

| | | Human classification | | | | |
|---|---|---|---|---|---|---|
| | | Accept | Reject - Population | Reject - Intervention | Reject - Study Design | Total |
| **Automatic classification** | Accept | 486 | 47 | 3 | 124 | 660 |
| | Reject - Population | 2 | 25 | 0 | 3 | 30 |
| | Reject - Intervention | 0 | 0 | 0 | 0 | 0 |
| | Reject - Study Design | 139 | 124 | 2 | 1454 | 1719 |
| | Total | 627 | 196 | 5 | 1581 | 2409* |

Abbreviations: AAM = advanced analytic methods; mCRPC = metastatic castration-resistant prostate cancer.
* AAM-2 was not able to classify 25 documents; therefore, the final sum is different from AAM-1

| Supplemental Table 12. AAM-2 (mCRPC): Confusion matrix for binary classification: Accept / Reject with Reason for Exclusion: Population | | | | | |
|---|---|---|---|---|---|
| Human classification | | | | | |
| Automatic classification | | Accept | Reject | Total | **Precision** |
| | Accept | 25 | 3 | 28 | 0.89 |
| | Reject | 124 | 1456 | 1580 | |
| | Total | 149 | 1459 | 1608 | |
| | | **Sensitivity** 0.17 | **Specificity** 1. | | |
| Abbreviations: AAM = advanced analytic methods; mNSCLC = metastatic non-small cell lung cancer. | | | | | |

| Supplemental Table 13. AAM-2 (mCRPC): Confusion matrix for binary classification: Accept / Reject with Reason for Exclusion: Intervention | | | | | |
|---|---|---|---|---|---|
| Human classification | | | | | |
| Automatic classification | | Accept | Reject | Total | **Precision** |
| | Accept | 0 | 0 | 0 | n/a |
| | Reject | 2 | 1606 | 1608 | |
| | Total | 2 | 1606 | 1608 | |
| | | **Sensitivity** 0. | **Specificity** 1. | | |
| Abbreviations: AAM = advanced analytic methods; mNSCLC = metastatic non-small cell lung cancer. | | | | | |

**Supplemental Table 14. AAM-2 (mCRPC): Confusion matrix for binary classification: Accept / Reject with Reason for Exclusion: Study Design**

| | | Human classification | | | |
|---|---|---|---|---|---|
| | | Accept | Reject | Total | **Precision** |
| **Automatic classification** | Accept | 1454 | 126 | 1580 | 0.92 |
| | Reject | 3 | 25 | 28 | |
| | Total | 1457 | 151 | 1608 | |
| | | **Sensitivity** 1. | **Specificity** 0.17 | | |

Abbreviations: AAM = advanced analytic methods; mNSCLC = metastatic non-small cell lung cancer.
